# Potential modes of COVID-19 transmission from human eye revealed by single-cell atlas

**DOI:** 10.1101/2020.05.09.085613

**Authors:** Kiyofumi Hamashima, Pradeep Gautam, Katherine Anne Lau, Chan Woon Khiong, Timothy A Blenkinsop, Hu Li, Yuin-Han Loh

**Author notes:** These authors contributed equally to this work (Kiyofumi Hamashima, Pradeep Gautam). Correspondence: Timothy A Blenkinsop, Icahn School of Medicine at Mount Sinai, New York, NY 10029, USA, Hu Li, Center for Individualized Medicine, Department of Molecular Pharmacology & Experimental Therapeutics, Mayo Clinic, Rochester, Minnesota 55905, USA, Yuin-Han Loh, Epigenetics and Cell Fates Laboratory, A*STAR, Institute of Molecular and Cell Biology, 61 Biopolis Drive Proteos, Singapore 138673, Singapore.

## Abstract

There is a pressing urgency to understand the entry route of SARS-CoV-2 viruses into the human body. SARS-CoV-2 viruses enter through ACE2 receptors after the S proteins of the virus are primed by proteases such as TMPRSS2. Most studies focused on the airway epithelial and lung alveolar cells as the route of infection, while the mode of transmission through the ocular route is not well established. Here, we profiled the presence of SARS-CoV-2 receptors and receptor-associated enzymes at single-cell resolution of thirty-three human ocular cell types. We identified unique populations of corneal cells with high ACE2 expression, among which the conjunctival cells co-expressed both ACE2 and TMPRSS2, suggesting that they could serve as the entry points for the virus. Integrative analysis further models the signaling and transcription regulon networks involved in the infection of distinct corneal cells. Our work constitutes a unique resource for the development of new treatments and management of COVID-19.

## Introduction

In November 2019, a novel coronavirus emerged in Wuhan, Hubei province, China, causing an outbreak of the infectious respiratory disease now known as COVID-19. As of 1 May 2020, the number of cases worldwide has exceeded 3 million, with over 233,000 deaths. COVID-19 is caused by the virus known as SARS-CoV-2, which utilizes angiotensin-converting enzyme 2 (ACE2) as a receptor for entry into host cells [1, 2]. Recently, a study by Wang et al. also reported Basigin (BSG), also known as CD147, as a novel receptor for SARS-CoV-2 entry into host cells [3]. The serum protease transmembrane protease serine 2 (TMPRSS2) is a key player in cleaving and activating the spike (S) protein of SARS-CoV-2 for viral entry, although the cysteine proteases cathepsin B and L (CTSB/CTSL) are also able to perform such a function to a smaller extent [1]. By analysing the expression of these genes across multiple tissues, we can deduce the cell types that could potentially be infected by SARS-CoV-2.

Studies analysing the gene and protein expression of ACE2 and TMPRSS2 across various tissues and organs have identified the organs at risk during SARS-CoV-2 infection and have also located specific cell types that may serve as target cells for infection of SARS-CoV-2[4, 5]. These organs include the heart, lungs, kidneys and ileum, while cell types vulnerable to infection include type II alveolar cells, myocardial cells and ileum epithelial cells. With BSG as a novel receptor for SARS-CoV-2 entry into host cells along with successful infection of MT-2 and A3.01 cell lines [6], this risk may extend to T lymphocytes as well which are known to express BSG on their cell surface. However, the distribution of BSG across the various tissues and its possible correlation with SARS-CoV-2 infection has not been well documented yet. Notably, the cornea and conjunctiva were also found to be potential targets for infection of SARS-CoV-2, suggesting a possible role for the eye in COVID-19 [5]. Overall, all these findings provide a clearer insight into the pathology of COVID-19 and potential novel modes of transmission.

Current evidence suggests that COVID-19 is primarily spread via respiratory droplets produced by infected individuals and contact routes [7]. However, other modes of transmission are also emerging as more and more studies report novel routes of viral invasion. One such mode of transmission is the ocular route, which has been implicated in recent studies and anecdotal reports [8-10]. Earlier studies also indicate that SARS-CoV, which is closely related to SARS-CoV-2 [11], is able to enter the body via indirect or direct contact with mucosal membranes in the eyes, suggesting that SARS-CoV-2 may also have the ability to gain entry via the ocular tissues [12, 13]. Recently, a study by Hui et al. observed that SARS-CoV-2 was able to infect conjunctival cells ex vivo and undergo productive replication, implying that the ocular route may emerge as a significant mode of transmission of COVID-19. [14].

Although the receptors used by SARS-CoV-2 to infect ocular cells have not been characterised, previous studies have pointed to possible roles for ACE2 and TMPRSS2 for SARS-CoV-2 entry in ocular cells. Analysis of single-cell RNA-sequencing (scRNA-seq) datasets from tissues including those of the cornea, retina, brain and heart showed that tissues in the cornea also express ACE2 and that superficial conjunctival cells were found to co-express ACE2 and TMPRSS2 [5] to a relatively high extent. In a separate study, immunolocalization of BSG in the human eye revealed BSG expression in certain ocular tissues including those of the corneal and conjunctival epithelium [15], indicating that BSG could also serve as a receptor for SARS-CoV-2 entry into certain ocular cells. It is also reported that SARS-CoV-2 could be present in ocular fluids and cause inflammation of the conjunctiva [16, 17]. Viral entry via the ocular route could result in subsequent manifestation of respiratory disease via the nasolacrimal system [12]. However, it remains unclear whether this route is a major mode of transmission of COVID-19 and which receptors are used by SARS-CoV-2 to infect ocular cells. Thus, more detailed characterisation of the mechanism by which SARS-CoV-2 infects ocular cells is urgently needed.

In this study, we use scRNA-seq datasets to analyse the expression of ACE2 and other SARS-CoV-2 entry genes, including BSG, TMPRSS2, cathepsin B, and cathepsin L, in ocular tissues to determine the mechanism by which SARS-CoV-2 infects ocular cells. Should the ocular route prove to be a significant mechanism of transmission, these findings will have implications for designing or selecting vaccines and antivirals that effectively block the ocular route as well.

## Results

### Identification of ACE2+ cells in Human Ocular Landscape

We prepared a single-cell atlas of different cell types from tissues of the human eye (Fig 1a). A high-resolution transcriptional landscape of 33,445 single-cells was mapped (Gautam and Hamashima et. al, manuscript under preparation). Following that, we inspected gene expression across tissues and thirty-three cell types in the human eye, discovering that ACE2 expression remains almost exclusive to a small percentage of cells in the cornea. In addition, within the cornea, at-least 6% of cells have high ACE2 gene expression compared to the rest of the ocular tissues (Fig 1c). We observed that conjunctival cells co-expressed ACE2 and TMPRSS2 which was also reported by Sungnak et al. [4]. Ocular epithelial cells also showed ACE2 expression, although that of TMPRSS2 was not detected. However, epithelial cells expressed genes coding for the cysteine proteases CTSB and CTSL (Fig 1c) which are also able to prime the SARS-CoV-2 S1 protein, suggesting they may still be vulnerable to viral infection despite having no observable TMPRSS2 expression. Besides ACE2, we analysed the expression of BSG (CD147) among the human eye cell types as it could be another receptor/co-receptor for SARS-COV-2 entry into host cells [9]. However, with BSG we did not observe any specificity in cell types. We also checked for expression of receptors or entry-related enzymes specific for other viruses such as the coronaviruses HCoV-229E and MERS-CoV along with influenza viruses. Besides influenza viruses, we could not find the expression of entry-associated genes specific to other viruses. This is consistent with previous reports that MERS-CoV infection does not show any ocular complications [10]. The presence of influenza receptors in the cornea is consistent with reports that this tissue is involved in subsequent manifestation of respiratory disease via the nasolacrimal route [5].

**Figure 1.**
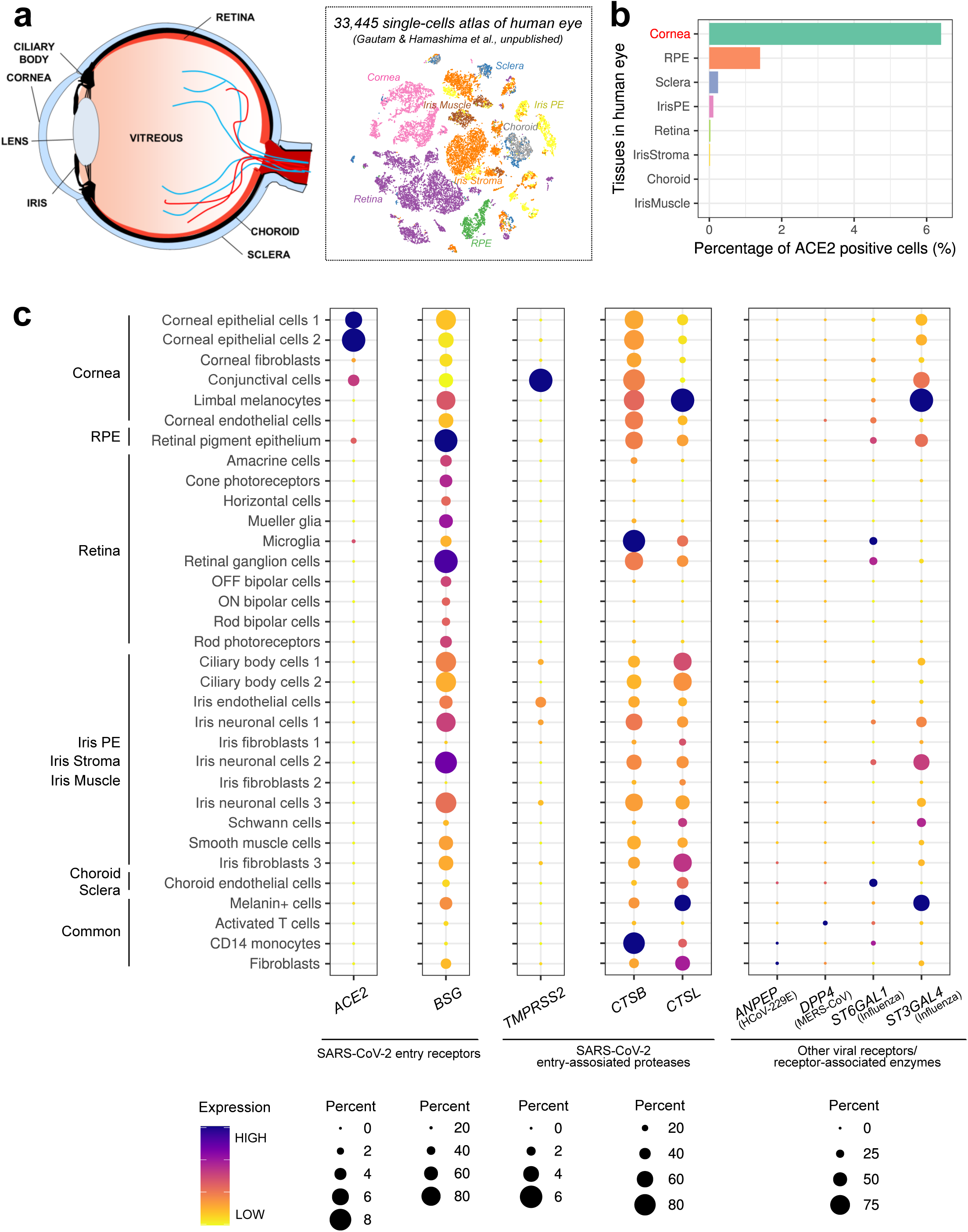
SARS-CoV-2 receptor *ACE2* is predominantly expressed in corneal epithelial cells in the human eye. **a**. Highlighted region of human eye selected for single-cell analyses (left panel) and the obtained human eye atlas visualized on t-SNE plot (right panel). 33,445 single-cells transcriptional landscape has been mapped at high resolution (Gautam and Hamashima et al., manuscript in preparation). **b**. Barplot indicating the percentage of *ACE2*-expressing cells within each tissue in human eye. **c**. Bubble plots showing the level of expression of indicated virus entry-associated genes and the ratio of expressing cells in the indicated cell types. The color of each bubble represents the level of expression and the size represents the proportion of expressing cells.

### ACE2 correlation analysis reveals its partner genes in cornea

To elucidate the pathways that are activated after ACE2 binding with the SARS-CoV-2 S1 protein, we sought to understand the transcriptomic differences between ACE2+ cells and the rest of the cells in the cornea. Therefore, we searched for genes that have high correlation with ACE2 and TMPRSS2 and listed some of the top genes (Fig 2a). Gene Ontology (GO) analysis of this list showed that terms related to cell membrane dynamics are enriched in such cell types (Fig 2b). The significant GO terms were related to/involved in cadherin binding, cell-cell junction, membrane edge, golgi complex, endosomal complex. We zoomed into a few genes that contributed to the GO terms (Fig 2c,d,e). Firstly, cadherin binding genes were mostly upregulated in ACE2+ cells (Fig 2c), which was consistent with reports from analysis of the bronchoalveolar lavage fluid of COVID-19 patients [19]. Among these genes, the ATP-dependent RNA helicase DDX3X is predicted to be able to bind to the S protein of SARS-CoV-2[20] and CD46 is a receptor for the measles virus [21]. MICAL2 is found to be one of the regulators of influenza virus infection [22]. MLLT4 is one of the components which links nectins with F-actins in cellular connections. Previous studies have shown that several viruses utilize nectins in cellular junctions for transmission [23]. Therefore, from our analysis, it is evident that ACE2+ cells have many genes which may aid SARS-CoV-2 in infection and its replication cycle. Similarly, it is logical that endosomal/Golgi transport is upregulated in ACE2+ cells as coronavirus entry is mediated through the endo/lysosomal pathway in a proteolysis-dependent manner [24].

**Figure 2.**
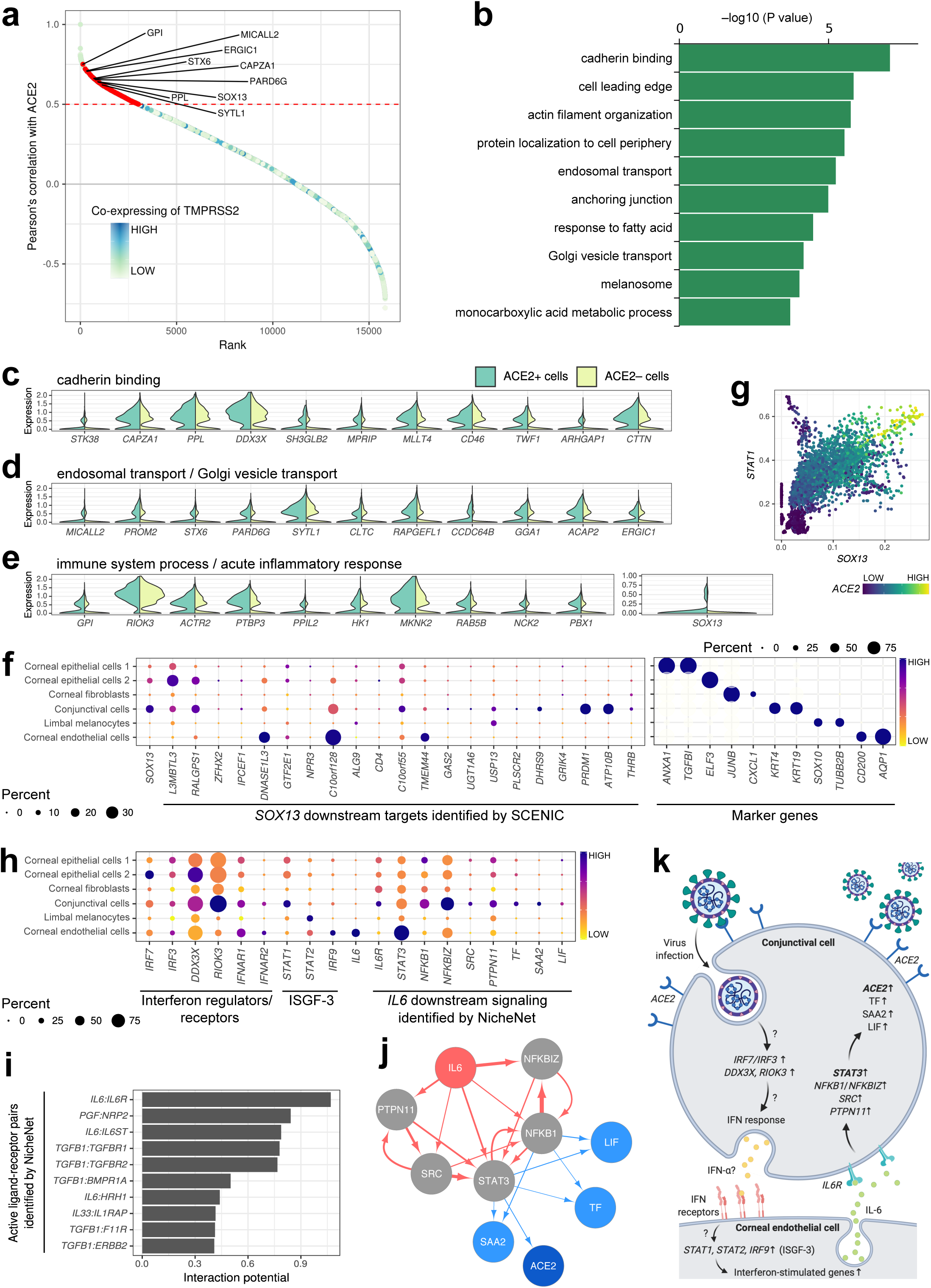
Single-cell atlas of corneal cells reveals possible pathways of virus transmission from the human eye. **a**. Scatter plot of Pearson’s correlation coefficients from association tests between *ACE2* and other individual genes in human corneal cells. Every gene is sorted by the correlation coefficient in a decreasing order. The color of each dot represents the level of co-expression of *TMPRSS2* with each gene (see the method section for details). The genes highly correlated with both *ACE2* (> 0.5 Pearson) and *TMPRSS2* are shown in red. 9 representative gene names (on the list of Fig. 2c-e) are labeled on the plot. **b**. Gene ontology (GO) analysis of genes that show high expression correlation with both *ACE2* and *TMPRSS2*. Top 10 enriched GO terms were listed. **c-e**. Violin plot showing expression of each GO term-associated gene in *ACE2* positive/negative cells in the cornea. From the gene-set used in GO analysis, highly expressing genes were selected for each GO term, cadherin binding (c), endosomal transport/Golgi vesicle transport (d) and immune system process/acute inflammatory response (e). The genes were ordered by the Pearson’s correlation coefficients shown in Fig. 2a. **f**. Bubble plots showing the level of expression of *SOX13* regulon (left panel) and marker genes (right panel), and the ratio of expressing cells in the indicated cell types in cornea. The *SOX13* regulon was computationally identified using SCENIC. **g**. Scatter plot indicating co-expression among *SOX13* (x-axis), *STAT1* (y-axis) and *ACE2* (color-scale) in a single-cell level. The expression level of each gene was imputed using MAGIC. **h**. Bubble plots showing the level of expression of interferon response-associated genes and the ratio of expressing cells in the indicated cell types in cornea. The *IL6* downstream genes were identified *in silico* by NicheNet analysis. **i**. Barplot indicating Nichenet-identified ligand-receptor interactions between corneal endothelial cells and conjunctival cells. Top 10 active pairs with high interaction potential (x-axis) are listed. **j**. Network showing high-confidence interactions forming putative signaling paths between the ligand *IL6* and its predicted target genes *TF, SAA2, LIF* and *ACE2*. The ligand node, target gene nodes and signaling/transcriptional regulator nodes are shown in red, blue and grey, respectively. Edge line thickness is proportional to the weight of the represented interaction in the weighted integrated networks. **k**. Proposed pathway of virus transmission via the route from *ACE2* expressed in conjunctival cells in the human eye.

Out of the highly correlated genes, SOX13 and PBX1 are the transcription factors that are active in ACE2+ cells. SOX13 is an islet cell autoantigen present in type I diabetic patients [25] and a lineage specification transcriptional factor for γd T cells which produce IL17 [26]. We checked the co-expression of SOX13 with STAT1 as STAT1 expression is highly correlated with ACE2 in COVID-19 patients and it acts upstream of ACE2 [27]. Subsequent MAGIC imputation revealed that cells that have high co-expression of SOX13 and STAT1 also have high expression of ACE2 (Fig 2g). Therefore, we wanted to identify genes that are downstream of SOX13 to elucidate a possible mechanism for SARS-CoV-2 infection of ocular cells following entry via ACE2. Transcription factors were identified by SCENIC analysis and their gene expression was checked across relevant ocular cell types (Fig 2f). We discovered that SOX13 expression is high in conjunctival cells, and some of the target genes of SOX13 (PLSCR2, PRDM1, ATP10B, DHRS9, THRB, and C10orf55) follow the same trend (Fig 2f). Genes like PLSCR2 have a role in fine tuning interferon expression after viral infection [28]. Additionally, PRDM1 directly targets STAT1 and affects interferon signalling pathways [29]. This shows that SOX13 may be one of the important transcription factors involved in SARS-CoV-2 infection of ACE2+ cells in the eye.

### Analysis of corneal signaling pathway after viral infection

The expression of interferon receptors (IFNAR1 and IFNAR2) in conjunctival and endothelial cells of cornea implies some sort of interaction between these cell types (Fig 2h). During SARS-CoV infection, the virus inhibits the expression of IRF3 and IRF7 transcription factors which are required for interferon induction [30]. DDX3X is known to be an interferon promoter [31] while RIOK3 is an adaptor protein required for IRF3 mediated interferon production [32]. We checked the expression of such proteins in all cell types of cornea and found out that DDX3X and RIOK3 are upregulated in conjunctival cells. Therefore, we checked for possible crosstalk occurring between endothelial cells and conjunctival cells, using NicheNet to find any significant ligand-receptor interactions between these two cell types (Fig 2i). Interestingly, we found IL6 gene expression to be high in corneal endothelial cells while ILF6R receptors were present in most of the conjunctival cells. It is established that IL6, along with other pro-inflammatory cytokines, are significantly elevated in COVID-19 patients and mostly seen in severe patients compared to non-severe patients [17]. We then proceeded to identify the genes that are activated by IL6 in conjunctival cells in the cornea.

By checking the expression of interferon regulators and receptors, we modelled the infection of conjunctival cells by viruses such as SARS-CoV-2 and crosstalk between conjunctival and endothelial cells. Following viral infection, interferon regulators are activated. Basal gene expression of these interferon regulators (IRF7, IRF3, DDX3, and RIOK3) in non-infected corneal cells shows that they have high expression in conjunctival cells. After infection, interferons released by infected cell types might communicate with endothelial cell types which have high expression of interferon receptors like IFNAR1 and IFNAR2 (Fig 2h). Interferon-stimulated genes (ISGs) such as STAT2 and IRF9 also show high expression in endothelial cells (Fig 2h). These factors likely stimulate the release of cytokines such as IL6 from endothelial cells, as shown by how IL6 expression was observed to be high in endothelial cells (Fig 2h). Further Nichenet analysis showed the genes downstream of IL6R and discovered STAT3, NFKB1, NFKBIZ, SRC, PTPN11, TF, SAA2, and LIF that had high regulatory potential (Fig 2h). These genes were also highly expressed in conjunctival cells (Fig 2h). IL6 has been shown to activate STAT3 transcription factors and the NF-KB pathway [33] while ACE2 has been shown to reduce the expression of STAT3 during lung inflammation [34]. Hence, there seems to be a tight balance between ACE2 and STAT3 expression. IL6 also enhances the uptake of transferrin in hepatocytes [35] and transferrin has been used by viruses to gain entry into host cells [36].

Using Nichenet data, we constructed a network depicting the genes that are activated by IL6 signalling in conjunctival cells and visualised this network using Cytoscape. Therefore, we present a model for conjunctival cells communication with endothelial cell types and reveal potential pathways for virus infection and response in ocular cell types. We propose that after infection of SARS-CoV-2 viruses, interferons produced by conjunctival cells and those interferons bind to interferon receptors in endothelial cells. In endothelial cells, interferon stimulated genes like IRF9 induce expression of IL6. IL6 then binds to IL6R receptors and induces the activation of the STAT3 based interferon pathway.

## Discussion

Using a single cell atlas of the human eye, we discovered that unique populations of cornea cells ACE2 expression. Further analysis revealed that these cells were mostly epithelial cell types present in the corneal and conjunctival cells. Conjunctival cells also showed expression of TMPRSS2 along with ACE2, suggesting that conjunctival sacs could be the entry point for SARS-COV-2 viruses. However, CTSB and CTSL were also expressed in ACE2+ corneal epithelial cell types indicating that SARS-CoV-2 could also infect certain population of epithelial cells in the cornea. While probing the genes that were highly co-expressed with ACE2 and TMPRSS2, we found SOX13 to be one of the interesting gene candidates. SOX13 is a transcriptional factor that affects the lineage specification of T cells and is also an autoantigen in Type 1 diabetes patients. The target genes of SOX13 such as PRDM1 and PLSCR2 affect interferon signalling pathways. Besides that, SOX13 also showed high coexpression with STAT1, which is one of the top predictors of ACE2 expression. This shows that SOX13 could be a viable drug target for modulating SARS-CoV-2 infection. While examining the microenvironment of corneal cell types, we detected high expression of IL6 in endothelial cell types. IL6 is one of the cytokines that is highly expressed in severe COVID-19 patients. This helped us to study the IL6 pathway in conjunctival cell types in the cornea. Using Nichenet analysis, we were able to find downstream genes of IL6 pathway and then construct a network for IL6 signalling in conjunctival cell types. With that we constructed a model of how conjunctival cells could be infected by viruses and their subsequent communication with endothelial cells. The endothelial cells release IL6 and bind to IL6R receptors in conjunctival cells. Together, that allowed us to determine the pathways that are potentially activated in response to IL6.

## Acknowledgements

We are grateful for Ankur Sharma for his helpful contributions to productive discussions. H.L. is supported by the Glenn Foundation for Medical Research, the Mayo Clinic Center for Biomedical Discovery and the Mayo Clinic Cancer Center. Y-H.L. is supported by the [NRF Investigatorship award -NRFI2018-02] and [JCO Development Programme Grant - 1534n00153] grants. Y-H.L. and L-F.Z. are supported by the Singapore National Research Foundation under its Cooperative Basic Research Grant administered by the Singapore Ministry of Health’s NationalMedical Research Council [NMRC/CBRG/0092/2015]. We are grateful to the Biomedical Research 565 Council, Agency for Science, Technology and Research, Singapore for research funding.

## Materials and methods

### Single-cell RNA-seq data processing and visualization

Single-cell RNA-seq of primary tissues in the human eye, followed by the generating high-resolution transcriptomic atlas was performed in our previous studies (Gautam and Hamashima et al., manuscript in preparation). Briefly, tissues (cornea, iris, ciliary body, neural retina, retinal pigmented epithelium, and choroid) from four individual post-mortem human adult eyes were dissociated by trypsin treatment. After freezing the cells, 10X Genomics single-cell RNA-seq was carried out according to the manufacturer’s instructions. Alignment to the reference genome, quantification, initial quality control (QC) and the downstream analyses were mainly performed using the 10X Genomics ‘cellranger’ pipeline (version 2.1.1 or 3.0.2) [37] and the R package ‘Seurat’ (version >3.0.1) [38] (https://github.com/satijalab/seurat). Total 33 cell classes were manually annotated based on the cluster-specific markers. The clustering result was visualized using t-distributed stochastic neighbour embedding (t-SNE). The generated Seurat object was exported and used in the present study. The expression profile of the selected genes was plotted on the bubble plot with the ‘DotPlot’ function in Seurat or violin plot with ‘geom_violin’ function in ggplot2 with the slight modification.

### Co-expression analysis of *ACE2* and *TMPRSS2*

The expression correlation between *ACE2* and other genes was drawn using Pearson’s correlation coefficient with gene expression value after MAGIC [39] (https://github.com/KrishnaswamyLab/MAGIC) imputation. MAGIC could restore missing values in the large sparse count matrix of single-cell RNA-seq. UMI count matrix of all 3,736 corneal cells was used for this analysis. We defined a gene with >0.5 Pearson as a co-expressing gene with *ACE2*. Due to the low *TMPRSS2* expression, the co-expression of *TMRPSS2* was analyzed as follows. First, individual cells were ordered by the imputed expression value for each gene and the population of cells in the top 10 percentile was extracted, gene by gene. Then, for each subset of cells, the number of cells expressing *TMPRSS2* was counted. We defined that when >=10 cells express *TMPRSS2* in each subset, the gene is co-expressed with *TMPRSS2* based on our prior analysis. The gene-set co-expressing with both *ACE2* and *TMPRSS2* was used in the gene-set enrichment analysis (GSEA) by Metascape [40] (http://metascape.org). Identification of *SOX13* downstream targets was performed using SCENIC [41] (https://github.com/aertslab/SCENIC) with default parameters. SCENIC is a powerful software which reconstructs a single-cell regulatory network based on the co-expression patterning between transcription factors (TFs) and potential target genes and the enrichment of the regulator’s binding motif.

### Cell-cell communication analysis

‘NicheNet’ [42] (https://github.com/saeyslab/nichenetr) was used to predict interaction molecules in the IL-6 downstream signaling in cornea. Among 3,736 corneal cells, 30 corneal endothelial cells and 159 conjunctival cells were selected as sender and receiver cells, respectively, in the analysis. Genes expressed in more than 5 % within each cell-type were retained. The gene-set having immune-associated GO terms (GO:0002376, immune system process or GO:0002526, acute inflammatory response) was set as target genes of interest. The ‘predict_ligand_activities’ function in the R package ‘nichenetr’ was run with default parameters. Predicted top 5 active ligand-receptor pairs were retained as highly potential interactions and top 3 potential targets of IL-6 signaling were featured in the figure. Based on the exported NicheNet data, the network graph was created using Cytoscape [43] (version 3.7.1) with a Compound Spring Embedder layout. The proposed pathway of virus transmission via the route from the human eye was illustrated using BioRender (https://biorender.com).

### Data availability

Single-cell RNA-seq data (human primary eyes) have been deposited in the Gene Expression Omnibus (GEO) under the accession code GSE147979, which will be released after publishing our previous studies (Gautam and Hamashima et al., manuscript in preparation).

### Code availability

All of the codes used in this study are available from the corresponding author upon reasonable request. Unless mentioned otherwise, all of the plots were generated using the R package ‘ggplot2’.

